# Activation of rod input in a model of retinal degeneration reverses retinal remodeling and induces formation of normal synapses, circuitry and visual signaling in the adult retina

**DOI:** 10.1101/469221

**Authors:** Tian Wang, Johan Pahlberg, Jon Cafaro, Alapakkam P. Sampath, Greg D. Field, Jeannie Chen

## Abstract

A major cause of human blindness is the death of rod photoreceptors. As rods degenerate, synaptic structures between rod and rod bipolar cells dissolve and the rod bipolar cells extend their dendrites and occasionally make aberrant contacts. Such changes are broadly observed in blinding disorders caused by photoreceptor cell death and is thought to occur in response to deafferentation. How the remodeled retinal circuit affect visual processing following rod rescue is not known. To address this question, we generated transgenic mice wherein a disrupted cGMP-gated channel (CNG) gene can be repaired at the endogenous locus and at different stages of degeneration by tamoxifen-inducible cre-mediated recombination. In normal rods, light-induced closure of CNG channels leads to hyperpolarization of the cell, reducing neurotransmitter release at the synapse. Similarly, rods lacking CNG channel exhibit a resting membrane potential that was ~10mV hyperpolarized compared to WT rods, indicating diminished glutamate release. Retinas from these mice undergo stereotypic retinal remodeling as a consequence of rod malfunction and degeneration. Upon tamoxifen-induced expression of CNG channels, rods recovered their structure and exhibited normal light responses. Moreover, we show that the adult mouse retina displays a surprising degree of plasticity upon activation of rod input. Wayward bipolar cell dendrites establish contact with rods to support normal synaptic transmission, which is propagated to the retinal ganglion cells. These findings demonstrate remarkable plasticity extending beyond the developmental period and support efforts to repair or replace defective rods in patients blinded by rod degeneration.

**Significance Statement:** Current strategies for treatment of neurodegenerative disorders are focused on the repair of the primary affected cell type. However, the defective neuron functions within a complex neural circuitry, which also becomes degraded during disease. It is not known whether a rescued neuron and the remodeled circuit will establish communication to regain normal function. We show that the adult mammalian neural retina exhibits a surprising degree of plasticity following rescue of rod photoreceptors. The wayward rod bipolar cell dendrites re-establish contact with rods to support normal synaptic transmission, which is propagated to the retinal ganglion cells. These findings support efforts to repair or replace defective rods in patients blinded by rod cell loss.

## Introduction

Diseases that afflict sensory systems typically result from deficiencies within the sensory receptor cells themselves, either within sensory transduction or synaptic transmission (Bermingham-McDonogh and Reh, 2011). Deficits in visual processing are no exception, with the majority of blinding diseases resulting from the dysfunction or death of the primary input cells, the retinal rod and cone photoreceptors (Quartilho et al., 2016). Synaptic remodeling of retinal circuits, in particular between photoreceptor cells and their downstream neurons, occur early in retinal degeneration (Soto and Kerschensteiner, 2015). Remodeling of bipolar and horizontal cell dendrites is thought to occur in response to deafferentation (Marc and Jones, 2003). Changes that occur include homeostatic down-regulation of synaptic structures, exuberant extension of dendritic processes which sometimes contact off-target sites (Marc and Jones, 2003; Puthussery and Taylor, 2010), and even switching of post-synaptic receptor types from mGluR to iGluR expression (Chua et al., 2009). In genetically inherited forms of retinal degeneration, synaptic changes may already occur during a critical period of retinal development. It is not known how these changes in retinal circuitry may ultimately limit recovery of normal vision, although several approaches are being implemented to rescue dying photoreceptors using gene therapy, or replace them with stem cells (Scholl et al., 2016; Garg et al., 2017; Yao et al., 2018). To address this gap in knowledge, this study focuses on cellular plasticity in retinal circuits of young adult mice with rod degeneration, and how the synaptic structures and circuits that receive rod input respond to rod rescue.

We genetically engineered a mouse line in which rod function can be uniformly rescued via tamoxifen-induced cre-mediated recombination. The line was generated to lack expression of the cyclic nucleotide gated (CNG) channel beta-1 subunit (CNGB1) due to an insertion of a neomycin cassette at the endogenous gene to disrupt expression (Wang and Chen, 2014; Wang et al., 2017a). This mouse model recapitulates the effects of mutations in human *CNGB1* and *CNGA1* genes that cause autosomal recessive retinitis pigmentosa (Biel and Michalakis, 2007). Without the CNGB1 subunit, the CNG channels in rod outer segments fail to form normally functioning channels, which leads to a slow form of rod death that occurs over 4-6 months (Zhang et al., 2009; Wang et al., 2017a), or longer (Hüttl et al., 2005). Importantly, the neomycin cassette is flanked by loxP sites, which allows for cre-mediated excision and the expression of CNGB1 from the endogenous locus. Thus, this mouse line provides an opportunity to introduce precisely a ‘cure’ for the underlying genetic defect at different time points during degeneration.

We use this novel mouse line to determine the extent to which activating rod input in the degenerating retina allows recovery of the structure and function of well-defined rod-driven retinal circuits in young adult mice. The lack of CNG channels caused stereotypic degenerative changes in the retina that included rhodopsin mislocalization, activation of Müller glia, and a reduction of pre- and post-synaptic proteins between rods and rod bipolar cells by as early as 4 weeks of age. Signal transmission from rods to rod bipolar cells was abrogated and sensitivity of retinal ganglion cells (RGCs) was reduced ~100-fold. Tamoxifen-induced restoration of CNG channel expression initiated at 4 weeks of age led to an expected recovery of rod photoreceptor function. Importantly, we show that initiation of rod input in the deafferented adult retina also induced a high degree of structural plasticity between rods and their primary postsynaptic partner, rod (ON) bipolar cells. Specifically, rod bipolar cell dendrites sprouted fine tips and mGluR6 clusters formed on these tips which made new synapses with rods. This structural transformation resulted in near-normal light responses in both bipolar cells and retinal ganglion cells, the output neurons of the retina. Our findings indicate substantial plasticity in the adult mammalian retina, suggesting favorable outcomes for interventions targeting the rescue of dysfunctional rods from death.

## Materials and Methods

### Generation of transgenic mice

The use of mice in these experiments was in accordance with the National Institutes of Health guidelines and the Institutional Animal Care and Use Committee of our respective universities. Targeting of the neoloxP to the *Cngb1* locus in mouse embryonic stem cells and generation of transgenic mice from verified stem cell clones were described previously (Chen et al., 2010). The CAGGCre-ER^TM^ transgenic line, Tg(CAG-cre/Esr1*)5Amc/J, was obtained from the Jackson Laboratory and crossed with *Cngb1^neo/neo^* mice.

### Tamoxifen treatment

One-hundred mg tamoxifen was dissolved in 500 µl 95% ethanol and diluted with 4.5 ml corn oil to give final concentration of 20 mg/ml. Four-week-old cre-positive were given a dose of 3 mg/25 g body weight by oral gavage for 4 or 7 consecutive days. In control experiments shown in Fig. 1, some cre-negative mice did not receive tamoxifen. For all other experiments, cre-negative littermate mice were also treated with tamoxifen to control for the possible effect of tamoxifen on photoreceptor cell survival (Wang et al., 2017b).

**Figure 1.**
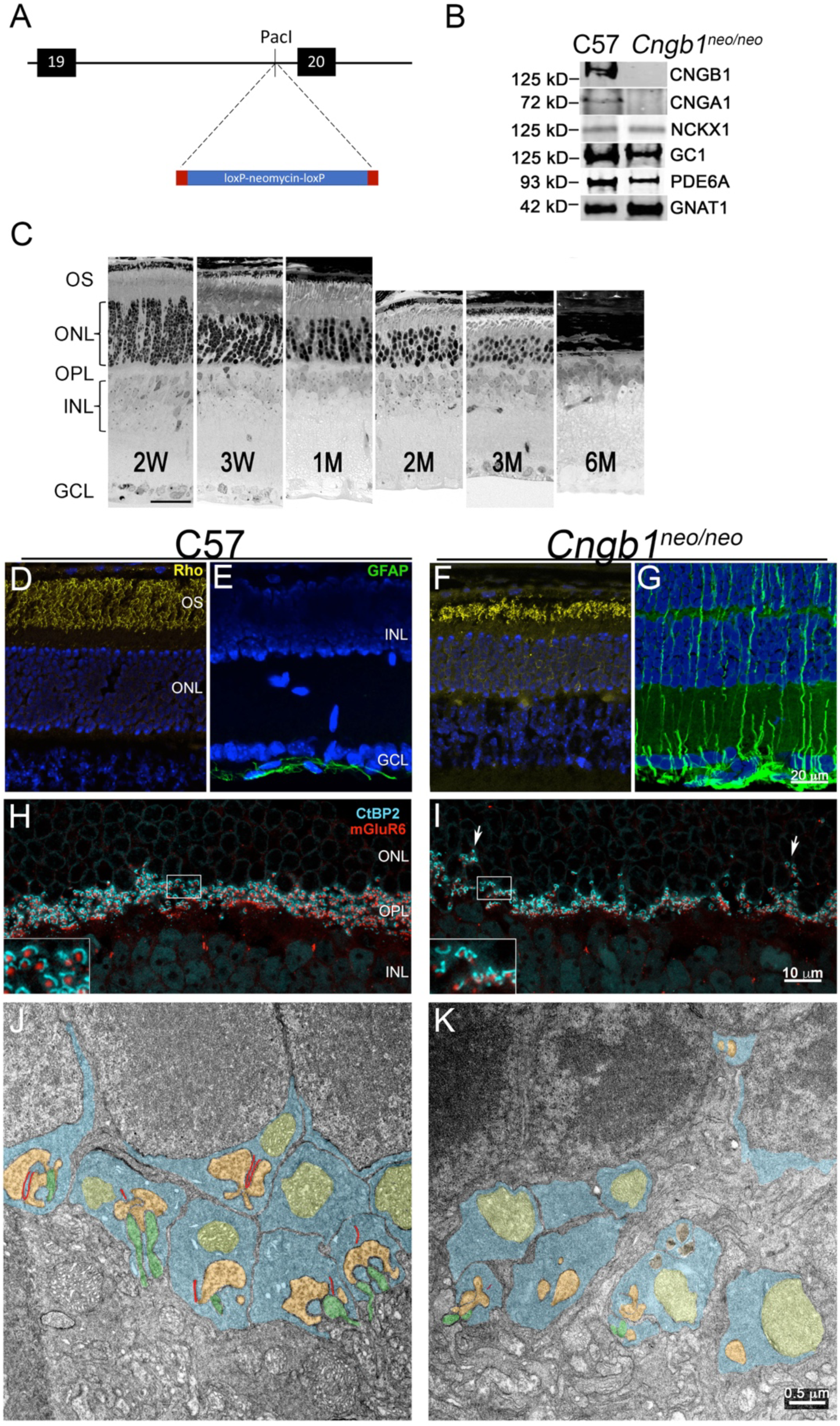
Retinas from *Cngb1^neo/neo^* mice exhibit stereotypic degenerative changes. (A) The 1.8 kb neomycin cassette, flanked by loxP sites, was inserted into intron 19 of the *Cngb1* gene. (B). Western blots of retinal homogenates from control and *Cngb1^neo/neo^* mice show that the neomycin insertion blocked expression of CNGB1, and down regulated expression of CNGA1 channel proteins. C. Light micrograph of representative retinal sections prepared from *Cngb1^neo/neo^* mice at the indicated ages. Scale bar = 20 µm. (D-I) Cryosections from 1 MO C57 (left panels) and 1 MO *Cngb1^neo/neo^* mice (right panels). Rhodopsin is localized to the outer segment (D) and GFAP to the inner limiting membrane proximal to the ganglion cell layer (E) in C57 retina. In contrast, the outer segment of mutant retina is shortened, and rhodopsin is mislocalized to the outer nuclear layer (F), and GFAP immunoreactivity extends to the entire retina (G). Nuclei are stained with DAPI (blue). (H) C57 and (I) *Cngb1^neo/neo^* retinal sections stained for synaptic ribbons (CtBP2, blue) of photoreceptors and mGluR6 puncta (red) of bipolar cell dendrites. Transmission electron microscopy of C57 (J) and *Cngb1^neo/neo^* (K) retinal sections. Color coding is as follows: rod spherule (blue), mitochondria (yellow), synaptic ribbon (red), horizontal cell (orange) and rod bipolar cell (green). OS, outer segment; ONL, outer nuclear layer; OPL, outer plexiform layer; INL, inner nuclear layer; GCL, ganglion cell layer.

### PCR genotyping

Genomic DNA was isolated from the neural retina. Three PCR primers were used to detect the presence or absence of the neoloxP cassette. Primer 1 sequence (GTTTTATGTAGCAGAGCAGGGAC) is located on intron 19, primer 2 sequence (GAGGAGTAGAAGGTGGCGC) is on neoloxP, and primer 3 sequence (CCACTCCTTAGTACATACCTAAGC) is located on exon 20. Product size of 620 bp from primer pairs (2+3) indicates the presence of neoloxP, and a 802 bp PCR band from primer pairs (1+3) indicates the absence of the neoloxP insert.

### Retinal morphology

Mice were rendered unconscious by isofluorane inhalation and immediately followed by cervical dislocation. Retinal sections were prepared as previously described (Concepcion and Chen, 2010; Wang and Chen, 2014). Briefly, before enucleation, eyes were marked for orientation by cauterization on the superior aspect of the cornea. Eyes were placed in ½ Karnovsky buffer (2.5% glutaraldehyde, 2% formaldehyde in 0.1 M cacodylate buffer, pH 7.2). The cornea and lens were removed, and the remaining eyecup was further fixed overnight. Fixed eyes were rinsed in 0.1 M cacodylate buffer, fixed for 1 h in 1% OsO_4_, dehydrated in graded EtOH and embedded in epoxy resin. Eyecups were hemi-sected along the superior-inferior axis, and one µm sections along the central meridian were obtained for light micrographs.

### Immunocytochemistry

Eyecups were prepared as described above, except the tissues were dissected in cold 4% formaldehyde in PBS and further fixed for 15 min on ice. For frozen sections, eyecups were rinsed in cold PBS, placed in 30% sucrose for 1 h, embedded in Tissue-Tek® O.C.T. Compound (Sakura® Finetek) and flash frozen in liquid N_2_. Ten µm frozen sections were obtained. For retinal flat mounts, four relaxing cuts (0°, 90°, 180°, 270°) were made on the edge of the neural retina and the flattened tissue was immobilized on a piece of nitrocellulose membrane (Whatman^®^, GE Healthcare Life Sciences), photoreceptor side down, as described (Anastassov et al., 2017). The tissues were incubated with the following antibodies: rhodopsin 1D4 (generously provided by R. Molday), GFAP (AB5804, Millipore), CtBP2 (612044, BD Biosciences), PKC (ab32376, Abcam), mGluR6 (generously provided by K. Martemyanov), ARR3 (generously provided by C. Craft). Images were acquired on a Zeiss LSM800 confocal microscope. For quantifications of mGluR6 puncta, images were imported into Fiji (ImageJ2), adjusted to similar threshold and the number and areas of puncta were quantified using the analyze particles function.

### Western blots

Each isolated retina was homogenized in 150 μl buffer (150mM NaCl, 50mM Tris pH 8.0, 0.1% NP-40, 0.5% deoxycholic acid, 0.1 mM PMSF and complete mini protease inhibitor (Roche Applied Sciences), incubated with DNase I (30U, Roche Applied Sciences) at room temperature for 30 min. An equal amount of retinal homogenate from each sample was electrophoresed on 4-12% Bis-Tris SDS-PAGE Gel (Invitrogen). Protein was transferred onto nitrocellulose membrane (Whatman^®^, GE Healthcare Life Sciences) and incubated overnight with the following primary antibodies: rabbit anti-PDE polyclonal antibody (PAB-06800, Cytosignal), rabbit anti-ROS-GC1 polyclonal antibody (sc50512, Santa Cruz), mouse Anti-G_t_α antibody (371740, EMD4Biosciences), rabbit polyclonal anti-GCAP1 and GCAP2 antibodies (Hoyo et al., 2014; Wang and Chen, 2014), mouse anti-CNGB1 4B1 antibody (Poetsch et al., 2001), mouse anti-CNGα antibody PMc 1D1 (Cook et al., 1989), mouse NCKX1 8H6 antibody (Vinberg et al., 2015) and mouse anti-Actin antibody (MAB1501, Millipore). The membranes were then incubated with fluorescently labeled secondary antibodies (1:10,000, LI-COR biosciences, 926-31081) at room temperature for 1 hour and detected by Odyssey infrared imaging system.

### Whole retina and single-cell recordings from rods and bipolar cells

Mice were maintained on a normal 12-hour day-night cycle and were dark-adapted overnight (>12-h) prior to experiments. All further manipulations were performed in total darkness under infrared illumination visualized with infrared image converters (BE Meyers, WA). Following euthanasia, eyes were enucleated, the lens and cornea were removed, and eyecups were stored in darkness at 32°C in Ames’ media buffered with sodium bicarbonate (Sigma, Cat# A1420) equilibrated with 5% CO_2_/ 95% O_2_.

*Trans*-retinal electroretinograms (ERGs) were recorded from isolated retinas as described previously (Pahlberg et al., 2017). Retinas were mounted photoreceptor side-up over a machined hole in a recording chamber. The tissue was superfused in darkness with 35-37°C Ames’ media buffered with sodium bicarbonate and equilibrated with 5% CO_2_/ 95% O_2_ (pH ~ 7.4). An additional 10 mM of BaCl was added to the solution facing the inner retina to mitigate Müller cell activity. The *trans*-retinal potential change to flashes of light, delivered from a standard light bench, was measured using Ag/AgCl half-cells connected to a differential amplifier (Model DP-311; Warner Instruments). Recordings were sampled at 1 kHz and low-pass filtered at 30 Hz.

Recordings of the photovoltage from individual rods and rod bipolar cells was made by whole-cell patch clamp from dark-adapted retinal slices as described previously (Pahlberg et al., 2017). Briefly, a small piece of dark-adapted retina was embedded in low-gelling temperature agar, slices were cut on a vibrating microtome, transferred into a recording chamber, and superfused with Ames’ media equilibrated with 5%CO_2_/95%O_2_ while maintained at 35-37°C. The pipette internal solution consisted of (in mM): 125 K-Aspartate, 10 KCl, 10 HEPES, 5 N-methyl glucamine-HEDTA, 0.5 CaCl_2_, 1 ATP-Mg, 0.2 GTP-Mg; pH was adjusted to 7.2 with N-methyl glucamine hydroxide. Light-evoked responses were recorded following the delivery of 10 ms flashes from a blue LED (λ_max_ ~ 470 nm, full width half maximum ~ 30 nm) whose strength varied from producing a just-measurable response, and increased by factors of 2. Recordings were sampled at 1 kHz and low-pass filtered at 300 Hz.

### Retinal ganglion cell recording, stimulation and analysis

Retinal ganglion cells were recorded from dorsal retina using a large scale, dense hexagonal multi-electrode array covering ~0.34mm^2^ of the retina (MEA, (Field et al., 2010) 519 electrodes with 30 μm spacing). The pigmented epithelium remained attached to the retina for these recordings. The retina was perfused with Ames’ solution (30-31°C) bubbled with 95/5% O2/CO_2_. Spikes were identified and assigned to specific RGCs on the MEA as previously described (Yu et al., 2017). Dim flashes were delivered at 3 s intervals using a 490 nm LED. Light intensity was controlled using pulse duration, 2-8 ms, and neutral density filters. Dim flash responses were measured by counting spikes on each trial within a 100 ms window that was centered on the peak of the peristimulus time histogram.

### Experimental design and statistical analyses

Because our initial studies did not show gender-specific differences, the genders were pooled. RGC response thresholds were measured from three *Cngb1ΔCaM* retinas (102-232 cells), five *Cngb1^neo/neo^* retinas (100-336 cells) and three *Cngb1^neo/neo^* rescue retinas (53-186 cells). Cumulitive threshold histograms were calculated in each tissue and averaged across all retinas within a condition. A two tailed Kolmogorov-Smirnov goodness-of-fit hypothesis test was used to assess the statistical difference between average cumulative histograms. The fraction of cells for which no response surpassed threshold was also measured in each recorded retina. A two sample t-test was used to evaluate significance between conditions.

## Results

### Generation of a novel animal model of genetically reversible rod degeneration

One challenge to identifying how plasticity among inner retinal neurons impacts functional recovery is the lack of an experimental system that is non-invasive and allows for stringent regulation of the timing and uniformity of rescue. For example, viral-mediated (gene therapy) approaches for treating rod dysfunction and death (1) take weeks for expression to occur; (2) they do not infect all targeted cells; (3) they may not drive proper protein expression levels; (4) and the subretinal injections used for viral delivery can damage the retina. A systematic investigation into the consequences of rod degeneration and subsequent rescue of the retinal circuitry requires an experimental system wherein both events occur uniformly in the retina. Towards this goal, a neoloxP cassette was inserted into intron 19 of the *Cngb1* gene by homologous recombination in mouse embryonic stem cells (Fig. 1A). Mice harboring this insertion were subsequently derived (*Cngb1^neo/neo^*). The presence of the cassette disrupted a splice site and prevented CNGB1 expression (Fig. 1B). Expression of CNGA1 was also substantially attenuated (Fig. 1B), a phenomenon attributed to mis-trafficking (Hüttl et al., 2005) and structural stability conferred by association of both subunits. The expression levels of other major phototransduction proteins were minimally perturbed in retinas of 1-month old (1 MO) mice (Fig. 1B, NCKX1, GC1, PDE6A, and GNAT1). Consistent with previous reports on conventional *Cngb1* knockout mice (Hüttl et al., 2005; Zhang et al., 2009), the lack of CNG channel expression led to a progressive thinning of the outer nuclear layer over the course of 6 months (Fig. 1C). At two-weeks, the outer nuclear layer (ONL) containing primarily rod photoreceptor cell nuclei reached its maximum thickness. This thickness was reduced by ~20% in 1 MO mice and to ~50% in 2 MO mice. By 6 MO, the ONL was absent. Thus, these mice exhibit slow rod degeneration relative to other commonly used models of rod degenerative diseases, such as *rd1* (Farber and Lolley, 1974) and *rd10* mice (Chang et al., 2007).

As expected, the absence of the CNGB1 recapitulated the stereotypic sequence of events associated with rod degeneration (Marc and Jones, 2003; Puthussery and Taylor, 2010; Soto and Kerschensteiner, 2015). For example, rhodopsin mislocalization and activation of Müller glia were observed in 4-week old *Cngb1^neo/neo^* mice (Fig. 1, compare F, G with control retina, D and E).

We also observed in *Cngb1^neo/neo^* mice that synaptic contacts between rods and rod bipolar cells were structurally abnormal. Immunohistochemistry using a marker for the presynaptic ribbon protein CtBP2 (ribeye) and the post-synaptic glutamate receptor, mGluR6, revealed clear differences between *Cngb1^neo/neo^* and control retinas. In control retinas, these structures were closely apposed, and both were contained within a well-defined outer plexiform layer (Fig. 1H, OPL). However, in *Cngb1^neo/neo^* retinas these structures were more dispersed, with synaptic ribbons retracted from rod spherules and some were situated deep into the photoreceptor nuclear layer (ONL, Fig. 1I, arrows). The ultrastructure of rod synapses was further evaluated by transmission electron microscopy (TEM, Fig. 1J and K). A normal rod spherule (blue) contains a single large mitochondria (yellow) and encompasses a synaptic triad consisting of a single ribbon (red) along with horizontal (orange) and rod bipolar cell (green) processes. The majority of imaged rod spherules from control retinas exhibited this structure. Dyads were also frequently observed when one or another component was out of the plane of the TEM section. Of the 27 fields taken from 4 control C57 retinas, 122 synapses were counted (4.5 synapses per field) wherein 64% were triads and 36% were dyads. However, in 1 MO *Cngb1^neo/neo^* retinas, the frequency of observing synapses and triadic structures were both reduced (Fig. 1K). Of 31 TEM fields taken from 4 *Cngb1^neo/neo^* retinas, 64 synapses were counted (2.1 synapses per field) wherein only 22% were triads and 78% of were dyads. Note, at 1 MO, only 20% of rods had died, yet there was >50% reduction in contacts between rods and rod bipolar cells. Furthermore the contacts that persisted were largely abnormal in structure. These results demonstrate that *Cngb1^neo/neo^* mice exhibit stereotypic slow rod degeneration and that synaptic structures between rods and rod bipolar cells were disrupted.

### Lack of CNGB1 expression attenuated rod photoresponses and eliminated rod bipolar cell light responses

Previous work has indicated that lack of CNGB1 expression compromises rod vision (Biel and Michalakis, 2007). To verify compromised rod function in *Cngb1^neo/neo^* mice, we performed *ex vivo* whole-retina electroretinograms (ERG) under scotopic conditions. The ERG reflects the averaged activity across all retinal neurons (Granit, 1933). ERGs from C57 retinas exhibited a well characterized biphasic response (Fig 2A; see also (Saszik et al., 2002)) with the initial negative-voltage deflection (a-wave) indicative of the rod hyperpolarization to the flash stimulus, and the subsequent positive-voltage rebound indicative of predominantly the rod bipolar cell depolarization. Recordings were also performed on control *Cngb1∆CaM* mice in which the calmodulin binding site was removed (Chen et al., 2010). No differences were observed between *Cngb1∆CaM* and C57 retinas (data not shown).

ERGs from 1 MO *Cngb1^neo/neo^* retinas exhibit a diminished scotopic a-wave with reduced sensitivity (Fig. 2B), indicating minimal rod signaling without CNGB1 expression. The light response is not fully eliminated in these rods; this is likely due to residual activation of homomeric channels composed of CNGA1 (see Discussion). Furthermore, ERGs from *Cngb1^neo/neo^* retinas did not exhibit a b-wave under scotopic conditions, indicating a lack of measurable rod-to-rod bipolar signal transmission at this gross level. This observation complements the abnormal synaptic structures observed between rods and rod bipolar cells via light and electron microscopy. Together, these results indicate that synaptic transmission between rods and rod bipolar cells is severely dysfunctional in *Cngb1^neo/neo^* mice. Thus, we sought to determine the extent to which normal synaptic structures and transmission between rods and rod bipolar cells could be recovered by the rescue of CNGB1 expression in mature retinas.

**Figure 2.**
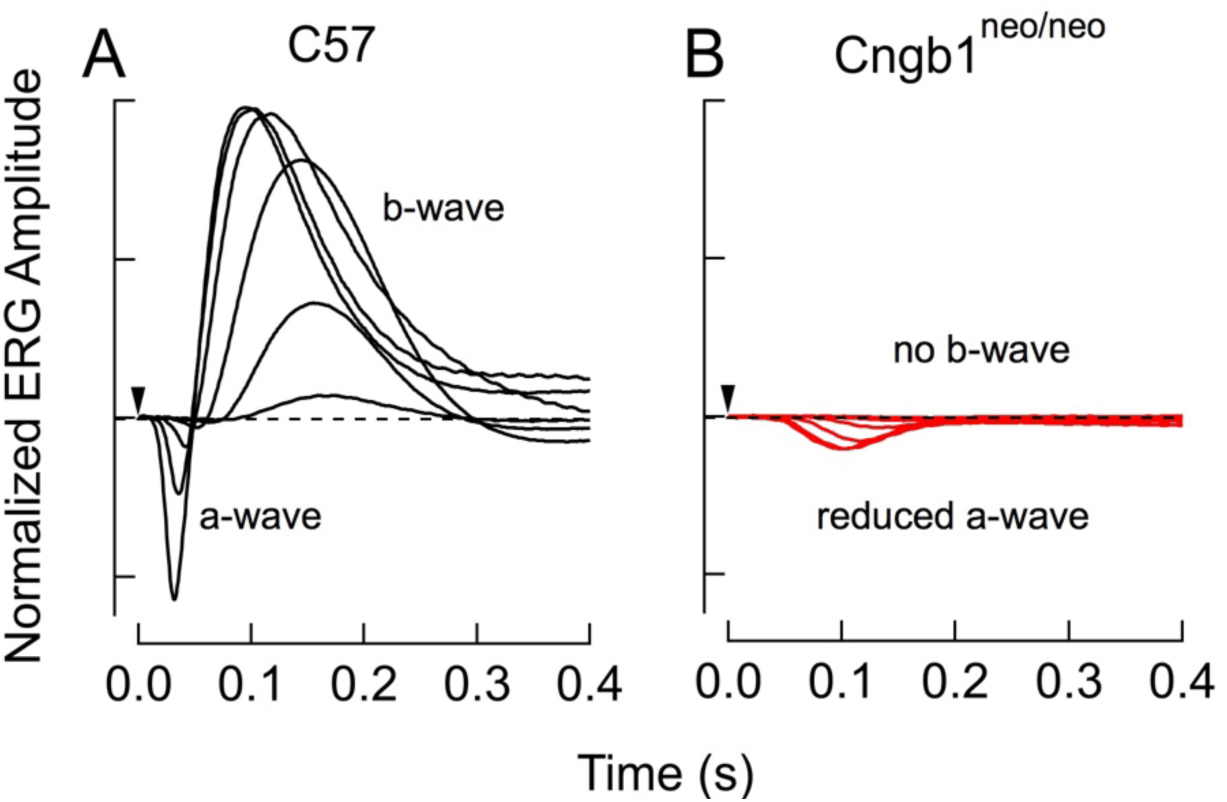
Characterization of rod function by *ex-vivo* electroretinography (ERGs). (A) ERG recordings from a 3 MO C57 mouse show normal a- and b-waves (rod and RBC responses, respectively). Flashes generated 2, 9, 35, 140, 560, and 2200 Rh*/rod (B) ERGs from 1 MO *Cngb1^neo/neo^* mice show total absence of the b-wave, consistent with disrupted rod-to-RBC signaling. Flashes generated 550, 2200, 8800, 18,000, and 35,000 Rh*/rod.

### Cre-mediated excision of the NeoLoxP cassette leads to normal CNGB1 expression

To activate CNGB1 expression in the the *Cngb1^neo/neo^* retina, we utilized the CAGGCre-ER^TM^ transgene (Hayashi and McMahon, 2002) to enable tamoxifen-dependent, cre-mediated excision of the NeoloxP cassette. We previously demonstrated that mice derived from germline excision of this cassette exhibit normal retinal morphology with a uniform and normal expression level of CNGB1 (Chen et al., 2010). The homologous recombination strategy that introduced the NeoloxP cassette also removed a stretch of 14 amino acids that encompassed the calmodulin binding domain on CNGB1 (Grunwald et al., 1998). Importantly, rods that expressed CNGB1ΔCaM exhibited normal light responses as mentioned above (see also (Chen et al., 2010)). We hypothesize that utilizing the CAGGCre-ER^TM^ transgene would provide temporal control over expression of the functional CNG channel. To determine the efficacy of cre-mediated excision of the neoloxP cassette, four-week-old cre-positive and cre-negative *Cngb1^neo/neo^* littermate mice were divided into two groups. One group was given tamoxifen for four-consecutive days by oral gavage, and the other group did not receive drug treatment.

A PCR strategy was designed to detect the extent of neoloxP excision in genomic DNA extracted from isolated retinas: the primer pair (2+3) detects the presence of the neoloxP insert, whereas primer pair (1+3) gives rise to a diagnostic band when the large neoloxP insert is excised (Fig. 3A). After four-consecutive days of tamoxifen treatment, both sets of primers produced positive bands. This result indicates a mixed population of cells at this stage, some of which have undergone excision while others have not. However, when tamoxifen treatment was given for seven-consecutive days a positive signal was detected only by primers (1+3). This result indicates that following a 7-day tamoxifen treatment, most, if not all cells have undergone neoloxP excision (Fig. 3A, bottom panels). Thus a 7-day treatment was used for further structural and functional studies.

**Figure 3.**
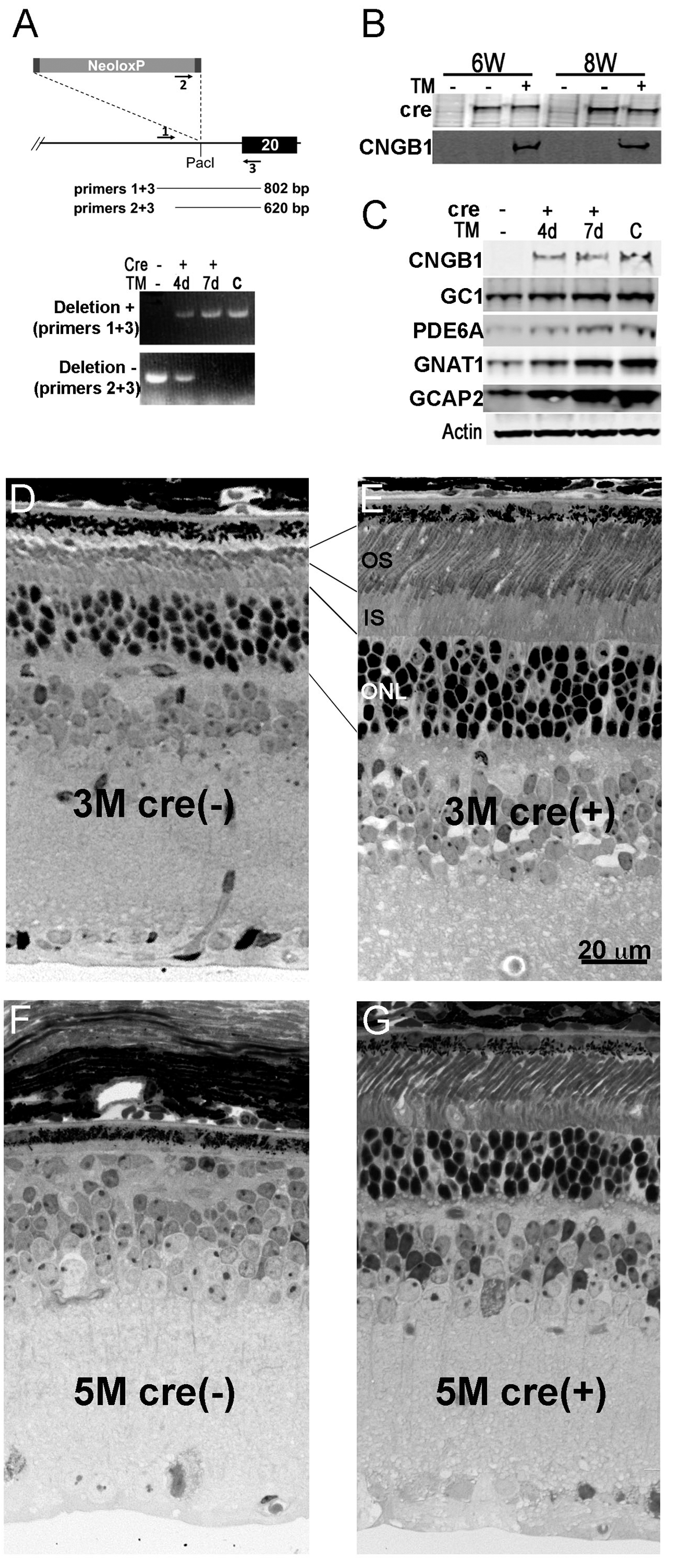
Excision of the floxed neomycin cassette restores CNG channel expression and rescues rod cell death. (A) PCR primers 1, 2, and 3 were designed to detect the presence or absence of the neoloxP cassette. Littermate mice were treated with tamoxifen (TM) for the indicated number of days starting at 4 weeks and retinal DNA was extracted from mice at 8 weeks. Control retinal DNA “c” is from a germline-floxed mouse wherein the neoloxP cassette has been removed in all tissues (*Cngb1*Δ*CaM*). (B) Western blots of retinal homogenates from cre-negative and cre-positive littermate mice of the indicated ages (6 weeks, 8 weeks) that were treated with TM or vehicle (corn oil) beginning at 4 weeks old for four consecutive days. (C) Western blot of retinal homogenate prepared from the contralateral eye from mice used in (A). Representative retinal morphology (N = 3) of 3 MO cre-negative (D) or cre-positive littermates (E) mice treated with tamoxifen for 7 consecutive days beginning at 4 weeks. TM-treatment of cre-positive mice showed improved outer segment (OS) length and thicker outer nuclear layer (ONL), indicating a halt on cell death. (F) cre-negative and (G) cre-positive littermates treated with tamoxifen for 4 consecutive days starting at P28, and retinal morphology was examined at 5 months of age (representative image from N ≥ 3). Scale bar = 20 µm.

To assess the level of protein expression at 6- or 8-weeks (corresponding to 1 or 3 weeks after drug treatment), Western blots were prepared from whole retinal homogenates from both cohorts (Fig. 3B). Expression of CNGB1 protein was observed only in tamoxifen-treated, cre-positive mice. No expression was observed in cre-positive mice without drug treatment, indicating a lack of basal recombinase activity. We next examined how excision of the neoloxP insert affected the expression of CNGB1 and other major phototransduction proteins. We found that following neoloxP excision, there was a striking increase in CNGB1 expression (Fig. 3C). There was also an increase in the detected levels of other phototransduction proteins GC1, PDE6A, GNAT1, and GCAP2 (Fig 3C). To determine if this is due to rod rescue, cre-negative and cre-positive *Cngb1^neo/neo^* littermate mice were administered tamoxifen for 7 consecutive days beginning at 4-weeks, and retinal sections were prepared from 3 MO mice (Fig. 3D and 3E). The ONL thickness was greater in cre-positive mice when compare to that from the cre-negative sibling mice, and the rod outer segment structure was organized and of normal length. Expression of CNGB1 exhibited a long term rescuing effect on rod survival (Fig. 3F and 3G, tamoxifen was administered for 4 consecutive days starting at P28), consistent with a previous report on AAV-mediated *Cngb1* gene replacement therapy (Koch et al., 2012). In sum, these data show that the *Cngb1^neo/neo^* mice allowed us to regulate the expression of CNGB1 from the endogenous locus in a temporally-controlled manner. Further, this excision is nearly complete with a 7-day tamoxifen treatment and that upon expression of CNGB1, the rods exhibit normal morphology and are stably rescued from cell death.

We measured the responsiveness of rod photoreceptors following drug treatment in patch-clamp recordings from individual rods in retinal slices. In voltage-clamp (V_m_ = -40 mV), rods from 1 MO *Cngb1^neo/neo^* mice displayed diminished response amplitudes (~6-fold) and a ~10-fold reduction in light sensitivity (Fig. 4A), a result consistent with the diminished a-wave in *ex vivo* ERG recordings (Fig. 2B). In current-clamp (*i* = 0), *Cngb1^neo/neo^* rods exhibit a resting membrane potential that was ~10 mV hyperpolarized compared to WT rods (-47 ± 1.3 mV (5) vs. -37 ± 2.3 mV (6), mean ± SEM). These results are consistent with reduced CNG channel expression (see Discussion) and indicate reduced glutamate release in darkness. However, rods from tamoxifen treated *Cngb1^neo/neo^* mice displayed responses with characteristics very similar to C57 mice (Fig. 4B), consistent with near-normal function and rescue of the photoreceptor layer (Fig. 3).

**Figure 4.**
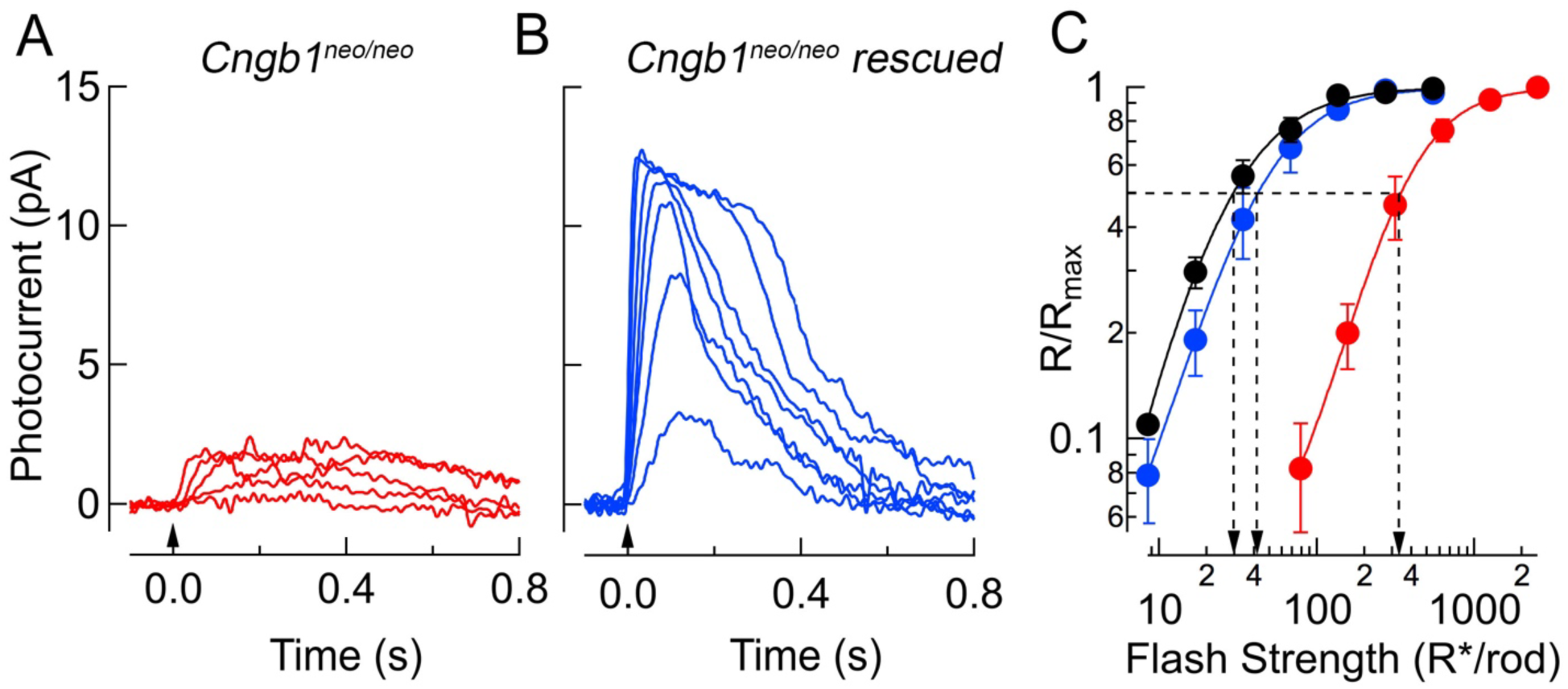
Light sensitivity is improved following expression of CNGB1. (A) Single cell recordings show small, desensitized response families in *Cngb1^neo/neo^* mice likely reflecting residual CNG channels composed of CNGA1 monomers. Flashes generated 79, 160, 310, 270, 1300, and 2500 Rh*/rod. (B) Following tamoxifen treatment, rod responses showed amplitudes and sensitivity resembling those of C57; flash strengths were 9, 17, 34, 68, 140, 270, and 540 Rh*/rod. (C) Response-intensity relationships from single-cell recordings display ~10-fold reduction in sensitivity between WT (black dots) and *Cngb1^neo/neo^* (red dots) (I_½_ values were 27 ± 4 (n=5) and 360 ± 8 (n=9), respectively). This sensitivity shift is nearly restored following reintroduction of the CNGB1 (blue dots; I_½_ = 43 ± 3 (n=10)).

### Expression of CNGB1 induces normal synaptic structures between rods and rod bipolar cells

Given that tamoxifen administration in *Cngb1^neo/neo^* mice rescued rods from death (Fig. 3E) and rescued normal rod light responses (Fig. 4), we next examined the synaptic contacts between rods and rod bipolar cells to determine how rod rescue impacts these structures. Tamoxifen treatment was initiated at 4-weeks for 7 consecutive days, and retinal structure was examined at 3M. Comparisons were made between 1 MO C57 and *Cngb1^neo/neo^* mice and 3 MO tamoxifen-treated mice to examine the effect of rod rescue that was initiated at 1M. Synaptic structures were labeled in retinal flat mounts stained for the presynaptic ribbon synapse protein (CtBP2, blue) and postsynaptic mGluR6 (orange, Fig. 5A, 5D and 5G). To distinguish between rod and cone synapses, cone pedicles were further labeled with the cone arrestin antibody (ARR3, green). Rod bipolar cell morphology, visualized by PKCa staining, and the mGluR6 puncta that decorate their dendritic tips are shown in retinal cross sections (Fig. 5B, 5E and 5H).

**Figure 5.**
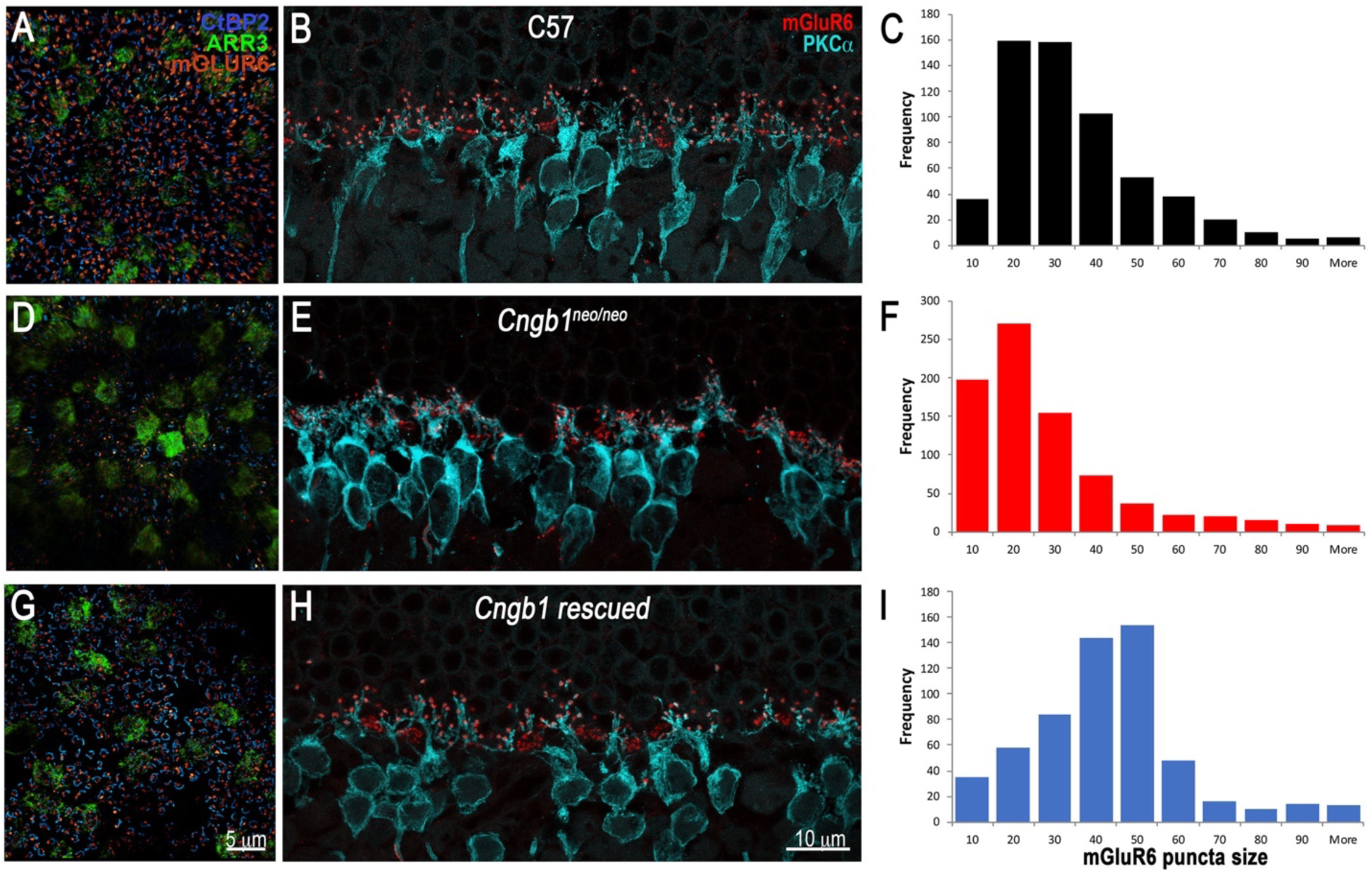
Expression of CNGB1 reverses pre- and postsynaptic retinal remodeling in *Cngb1^neo/neo^* retina. Shown are representative images from N>3 independent experiments. A, D and G are retinal flat mounts from 1 MO C57, 1 MO *Cngb1^neo/neo^* and 3 MO rescued *Cngb1* mice treated with tamoxifen for 7 consecutive days beginning at 4 weeks of age, respectively. The flat mounts were stained with the pre-synaptic ribbon marker, CtBP2 (blue), and post-synaptic marker mGluR6 (orange). Cone pedicles were visualized using cone arrestin, ARR3 (green). B, E and H are retinal cross sections from mice of the same genotype as the flat mounts. The retinal sections were stained with antibodies to mGluR6 (red) and the rod bipolar cell marker, PKCa (teal). C, F and I are frequency vs. mGluR6 puncta size distributions for 1 MO C57, 1 MO *Cngb1^neo/neo^* and 3 MO rescued *Cngb1* mice treated with tamoxifen.

Control retinas (C57), exhibited a close juxtaposition between the rod’s single ribbon and the mGluR6 puncta on the dendritic tips of rod bipolar cells (Fig. 5A and 5B; Fig. 1G). However, in *Cngb1^neo/neo^* retinas from 1 MO mice, both the number of synaptic ribbons and mGluR6 puncta were reduced (Fig. 5D). Rod bipolar cell dendrites were also unevenly distributed in *Cngb1^neo/neo^* retinas, and the size of the mGluR6 puncta appeared smaller and less uniform in shape (Fig. 5E and Fig. 1H). In contrast, retinas from the tamoxifen-treated, cre-positive littermates exhibited robust staining of synaptic ribbons along with their associated mGluR6 puncta (Fig. 5G). Furthermore, rod bipolar cell dendrites were evenly extended and the mGluR6 puncta were larger in size and appeared more uniform in shape (Fig. 5H). To quantify these changes, the number and size of mGluR6 puncta were measured (Fig. 5C, 5F and 5I). For the C57 retina, mGluR6 puncta size of 20- to 30-unit area were the most numerous, whereas in *Cngb1^neo/neo^* retinas smaller size puncta were more frequent (compare Fig. 5C and 5F). The distribution shifted back to larger puncta sizes in the tamoxifen-treated mice (Fig. 5I). These results indicate that inducing expression of CNGB1 in mature retina causes a recovery of synaptic structures between rods and rod bipolar cells.

### Rescue of CNGB1 expression in mature retina recovers rod bipolar light responses

The results above indicate a structural recovery of synapses following expression of CNGB1. In addition to this structural recovery, *ex vivo* whole-retina ERGs revealed a recovery of the rod bipolar cell-driven b-wave with amplitudes similar to control retinas (Fig. 6A). Thus, structural and functional measures broadly indicate recovery of synaptic function between rods and rod bipolar cells. To examine further synaptic function before and following rod rescue, we performed patch clamp recordings from rod bipolar cells in retinal slices. In untreated *Cngb1^neo/neo^* mice there was a complete absence of functional transmission of between rods and rod bipolar cells (Fig. 6C); 1 MO *Cngb1^neo/neo^* rod bipolar cells never yielded light-evoked responses (n=15 from 5 retinas), a result consistent with a lack of b-wave in *ex vivo* ERG recordings which represent mass-action of largely rod bipolar cells (Fig. 2B). However, in *Cngb1^neo/neo^* mice administered tamoxifen for 7 days at 4 weeks of age and recorded at 3 MO, rod bipolar cells exhibited robust light-evoked responses similar to control animals (Fig. 6D). The extent of functional recovery in rod bipolar cells was characterized in plots of the response amplitude versus the flash strength. These intensity-response relationships were fit with a Hill curve and compared quantitatively to control responses. The half-maximal flash strength (I_1/2_) increased by ~2-fold in rescued animals (Fig. 6E), consistent with some rod loss (see Discussion). However, other features of the rod bipolar light response that are critical for function near visual threshold had recovered to near control values. For example, the Hill exponent for the fit of the response-intensity relationship matched that in control, indicating a similar nonlinear relationship between the flash strength and the response amplitude. The extent of nonlinearity reflects the rate of glutamate release from rod synapses (Sampath and Rieke, 2004). In addition, the time course of rod bipolar cell responses was similar in rescued animals (Fig. 6A, 6C, 6E – dashed line), further indicating the anatomical and physiological recovery of synaptic transmission in tamoxifen-treated animals. These results indicate that rescuing rod function in the mature mouse retina produces a cascade of structural and functional recovery in synaptic transmission between rods and rod bipolar cells, and thus the primary rod pathway (Dacheux and Raviola, 1986).

**Figure 6.**
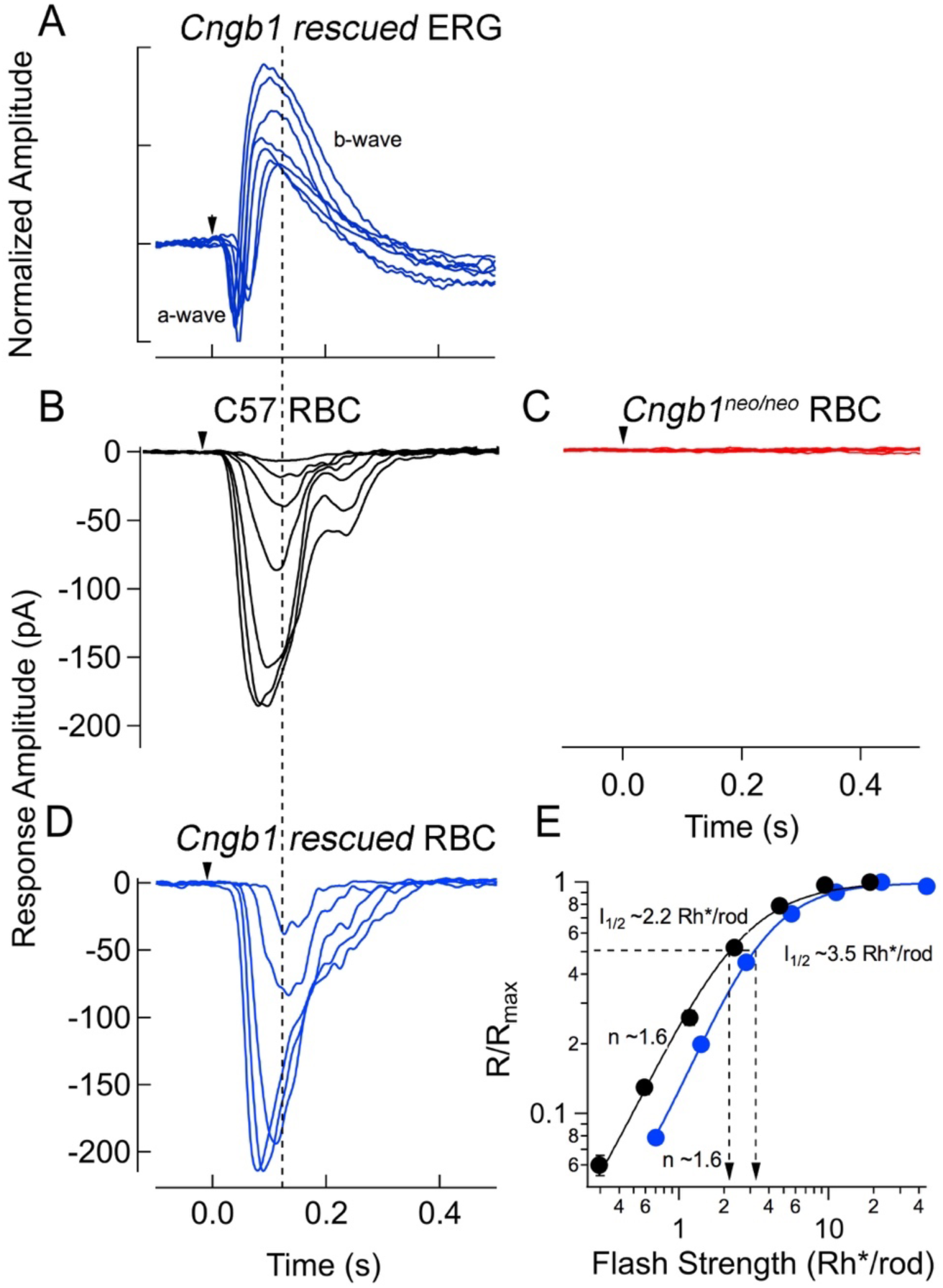
Physiological responses from rod bipolar cells in retinal slices. A. *Ex-vivo* ERG responses from 3 to 6 MO mice after tamoxifen treatments show near normal a- and b-waves. Flashes generated 2, 9, 35, 140, 550, and 2200 Rh*/rod. Please compare against Fig. 2A. Voltage-clamp rod bipolar cell recordings (V_m_ = -60 mV) from the following mice: B. C57 rod bipolar cells (2-3 MO); C. *Cngb1^neo/neo^* rod bipolar cells (1 MO); D. *Cngb1* tamoxifen-treated (3 MO). Flashes generated 2, 4, 8, 16, 31, 62 and 130 Rh*/rod for C57 rod bipolar cells, and 280, 560, 1100 and 2200 Rh*/rod for *Cngb1^neo/neo^* rod bipolar cells. Light-evoked responses were never observed in *Cngb1^neo/neo^* rod bipolar cells (15 cells across 5 retinas – flashes generated 2200 Rh*/rod). E.Response-intensity relationships from mean data show that this relationship is shifted to higher flash strengths in rescued mice, reflecting some rod loss. The Hill exponent of rod bipolar cells were similar to normal following rod recovery (Hill exponent = 1.6 ± 0.05 (n=12)), in support of a restoration of the normal rod-to-rod bipolar cell synaptic structure and the dark rate of glutamate release.

### Rescuing rods recovers absolute sensitivity of retinal output

Retinal ganglion cells (RGCs) provide the sole output from the retina and can integrate input from thousands of rods, making them the most light-sensitive cells in the retina (Chichilnisky and Rieke, 2005; Field and Sampath, 2017). RGC sensitivity relies on functioning photoreceptors and highly tuned synaptic connections via the primary rod pathway (Field and Rieke, 2002; Sampath and Rieke, 2004). To understand how rescuing rod function in the 1 MO *Cngb1^neo/neo^* retina impacts RGC sensitivity, we used a large-scale multi-electrode array (MEA) to record spikes from hundreds of RGCs. We tested the sensitivity of the RGCs by stimulating the retina with brief, dim flashes (0.001-10 Rh*/rod) and compared RGC responses in 3 MO control *Cngb1∆CaM*, *Cngb1^neo/neo^*, and 4-5 MO *Cngb1* tamoxifen rescued mice. Flashes producing less than 1 isomerization per rod faithfully produced spike rate modulations in many RGCs from control mice (Fig 7A shows an example cell). The same flash intensities did not reliably modulate the spike output of most RGCs in retinas from 3 MO old *Cngb1^neo/neo^* mice (Fig 7B shows an example cell). Indeed, most RGCs from *Cngb1^neo/neo^* mice did not show reliable responses until flash intensities exceeded 1 Rh*/rod (Fig 7B_3_). However, similar to the control retinas, RGC responses were often evident at low flash intensities in 4-5 MO *Cngb1* rescued mice (Fig 7C shows an example cell). These example cells suggest that rod and circuit functionality are broadly and stably rescued in some RGCs for *Cngb1* rescued mice.

To measure the extent that sensitivity across the RGC population recovered in *Cngb1* rescued mice, we quantified the response-threshold for all RGCs (N=1954) in MEA recordings from 11 mice (3 control *Cngb1ΔCaM* mice, 3 *Cngb1^neo/neo^* mice, 5 tamoxifen-treated *Cngb1* mice). RGC response thresholds were quantified as the lowest flash intensity needed to drive the average spike rate change two standard deviations above baseline (eg. Fig 7A_3_, 7B_3_, 7C_3_; see Methods). Average RGC response threshold distributions were similar between control and *Cngb1* rescued mice, but were significantly higher in untreated *Cngb1^neo/neo^* mice (Fig 7D; KS test, p<0.05). Additionally, the fraction of RGCs for which no-threshold response could be measured was similar between control and *Cngb1* rescued mice but significantly higher in untreated *Cngb1^neo/neo^* mice (Fig 7E; *t*-test, p<0.05). These results indicate a broad and lasting recovery of rod and circuit functions in *Cngb1* rescued mice.

### Discussion

In contrast to other neurons, rods and cones are depolarized in darkness and tonically release glutamate through ribbon synapses (Molday and Moritz, 2015), leading to saturation of rod-to-rod bipolar cell synapses (Sampath and Rieke, 2004). Light exposure causes graded hyperpolarization of the photoreceptor cell and suppression of glutamate release. Reductions in glutamate release from photoreceptors that occur during the early process of retinal degeneration lead to homeostatic changes in the downstream neurons and degrade the retinal circuit. This is seen in the dissolution of synaptic structures, dendritic sprouting, formation of ectopic contacts, and gliosis (Marc et al., 2003; Puthussery and Taylor, 2010). Although strategies to rescue and restore function in defective photoreceptors have shown success for regaining some visual function, gene and stem cell therapies for visual restoration are often implemented in the adult; how well these rescued neurons reinstate their detailed circuitries in the remodeled retina is not known. Here we examined functional restoration at the level of inner retinal cells and defined rod-driven circuits in the young adult mouse retina. We show that repairing a primary genetic defect in rods not only restored rod function, but also recovered normal synaptic connectivity with remodeled second order rod bipolar cells.

### Rod-to-rod bipolar cell synaptic contacts are reduced in *Cngb1^neo/neo^* retina and do not support synaptic transmission

The relatively slow photoreceptor degeneration we observe in the *Cngb1^neo/neo^* mouse model, and observed in human patients (Bareil et al., 2001; Biel and Michalakis, 2007), may be due to the fact that it is not a functional null. A small but measurable light response persisted in *Cngb1^neo/neo^* rods from 1 MO mice (Fig. 4B). The residual light response is probably due to the presence of homomeric channels composed of CNGA1 subunits, which are capable of mediating a diminished and desensitized cGMP-dependent current (Kaupp et al., 1989). The small current would reduce Ca^2+^ influx to the outer segment, causing increased levels of cGMP through stimulation of guanylyl cyclases by Ca^2+^-free guanylyl cyclase activating proteins 1 and 2 (Mendez et al., 2001; Dizhoor et al., 2010). Elevated cGMP has been shown to be a driver of rod degeneration through activation of protein kinase G (Ma et al., 2015; Wang et al., 2017a).

Despite the diminished rod light responses in the *Cngb1^neo/neo^* mice, the utter lack of evidence for light-evoked transmission between rods and rod bipolar cells was unexpected, especially given our ultrastructural evidence showing that the number of triads in the *Cngb1^neo/neo^* rod spherules was reduced, but not fully eliminated (Fig. 1). Therefore, it is surprising that ERGs from *Cngb1^neo/neo^* mice did not exhibit a scotopic b-wave, which is largely contributed by rod bipolar cells (Fig. 2B; (Saszik et al., 2002). This result was further corroborated by patch-clamp measurements from rod bipolar cells that revealed no light response, even for bright flashes delivering ~2000 Rh*/rod (Fig. 6C). We speculate that this defect in synaptic transmission is due to diminished glutamate release at the ribbon synapse given that the lack of CNG channels should act as a source of “equivalent light”, similar to light-induced closure of CNG channels (see also Sampath and Rieke, 2004; Dunn et al., 2006). Supporting this idea, the resting membrane potential of *Cngb1^neo/neo^* rods are ~10 mV hyperpolarized due to their smaller dark current (Fig. 4A). At the rod’s normal resting potential in darkness (~40 mV), calcium enters through the voltage gated channel (Ca_v_1.4) and supports tonic glutamate release at the ribbon synapse (Waldner et al., 2018). Thus the hyperpolarizing shift in resting potential of *Cngb1^neo/neo^* rods predicts attenuated glutamate release from the rod spherule.

Interestingly, suppression of glutamate release at the rod synapse is strongly correlated with synaptic remodeling. Examples include blockade of glutamate release by tetanus toxin (Cao et al., 2015), in knockout mice that lack the presynaptic Ca_V_1.4 Ca^2+^ channel (Mansergh et al., 2005), and in human patients diagnosed with congenital stationary night blindness (CSNB2) that harbor null mutations in the gene encoding Ca_V_1.4 (Bech-Hansen et al., 1998; Boycott et al., 2000). Calcium entry through Ca_V_1.4 channel is required for neurotransmitter release at the ribbon synapse of both rods and cones. The absence of CaBP4 (Haeseleer et al., 2004) or a2d4 (Wang et al., 2017c) that bind and regulate the activity of Ca_V_1.4, also manifest in retinal remodeling in knockout mice. These plastic changes occurred with minimal photoreceptor cell loss, suggesting that synaptic remodeling is likely driven by suppression of neural transmission, or deafferentation, rather than photoreceptor cell death *per se*. Modest synaptic changes were also observed in RIBEYE knockout retinas. RIBEYE is an essential component of the synaptic ribbon, and its absence abolished all presynaptic ribbons in the retina and severely impaired fast and sustained neurotransmitter release (Maxeiner et al., 2016). Spontaneous miniature release continues to occur without the synaptic ribbon, which may explain the milder retinal remodeling phenotype observed in the RIBEYE knockout retina (Maxeiner et al., 2016).

Previous studies on two independent lines of *Cngb1^-/-^* mice have similarly reported a substantial reduction of light responses from *Cngb1^-/-^* rods, but in both mouse lines the presence of a rod bipolar cell driven b-wave was observed (Hüttl et al., 2005; Zhang et al., 2009). The reason behind the discrepancy between those and our results is not clear, but may be due to differences in: 1) the degree of rod hyperpolarization caused by the number of functioning homomeric CNGA1 channels and hence the amount of glutamate released by rods in the different mouse lines, 2) the *in vivo* vs. *ex vivo* ERG measurements, or 3) mouse genetic backgrounds.

### Adult rod bipolar cells demonstrate plastic changes to establish functional contacts with rescued rods

The developmental time window for the formation of the rod to rod bipolar cell synapse in mice appears to be from eye opening to postnatal day 30 (P30), during which synaptic proteins are expressed, pre- and post-synaptic molecular complexes form, and the rod bipolar cells develop the appropriate number of dendritic tips that make synaptic contacts with rods (Anastassov et al., 2017). Some of the molecules that guide neurite growth during development are absent at maturity (D’Orazi et al., 2014), and if functional connectivity of the neural retina can only occur during a critical period in development, then one would expect that the adult retina may lack the ability to make such connections when rod activity is switched on after this time window. Such developmental processes would have been disrupted in *Cngb1^neo/neo^* retinas, wherein pronounced retinal remodeling is evident by P30 (Fig. 1). We show that tamoxifen-induced CNGB1 expression between P28-P34 led to establishment of the rod’s circulating current in darkness and normal light responses (Fig. 4). Concomittantly, structural changes were observed at the synapse: rod bipolar cells elaborated fine dendritic tips, mGluR6 receptors clustered on these tips which came in close contact with presynaptic ribbons (Fig. 5G-5I). These newly formed synaptic structures supported normal neural transmission, as shown by ERG recordings and patch recordings from rod bipolar cells (Fig. 6), and light sensitivity in increased in RGCs (Fig. 7). We hypothesize that these changes may be initiated by glutamate release at the rod’s synapse, similar to that which occurs at the cortex, where focal uncaging of glutamate in mouse cortical layer 2/3 pyramidal neurons triggered spinogenesis from the dendrite shaft in a location-specific manner (Kwon and Sabatini, 2011).

**Figure 7.**
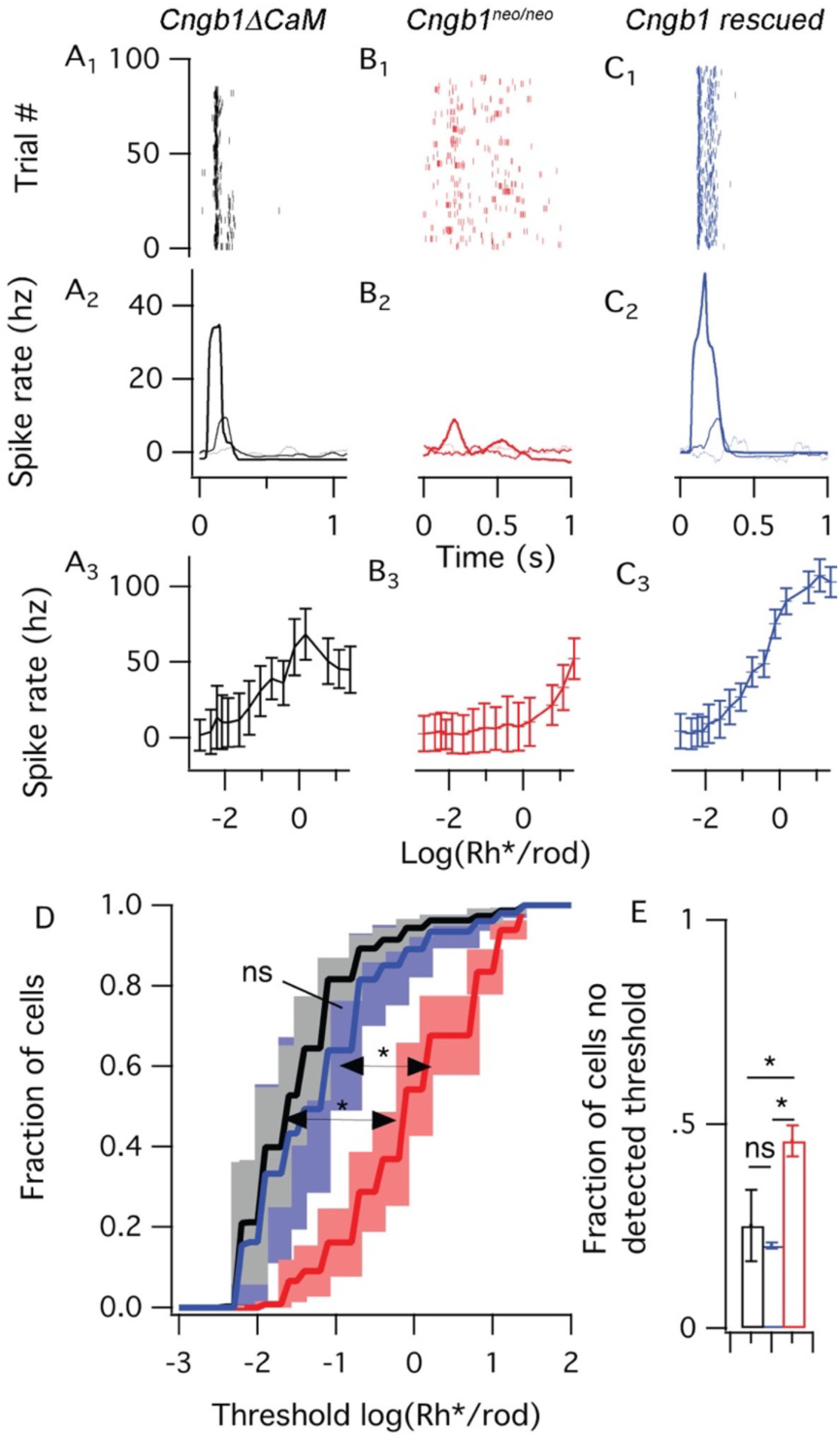
Dim flash responses from retinal ganglion cells (RGCs) in whole mount retina show recovery from early rod rescue. The top panels (A-C_1_) show spike times of 100 trials of 3 example cells to a single dim flash (0.75 Rh*/rod). The middle panels (A-C_2_) shows PSTHs for three increasingly bright dim flashes (0.002, 0.02, 0.75, Rh*/rod). The bottom panels (A-C_3_) show the mean spike rate ± SD, measured on each trial in a 100 ms window around the peak of the PSTH. Dim flash thresholds were estimated from these curves for 1954 cells. The average cumulative distribution function show higher thresholds responses in *Cngb1^neo/neo^* mice RGCs than *Cngb1∆CaM* and tamoxifen-treated *Cngb1^neo/neo^* mice RGCs (D). The shaded regions illustrating SEMs across (3-5 experimental preparations). Additionally, flash thresholds could not be identified in a larger portion of RGCs from *Cngb1^neo/neo^* mice (E).

Plasticity at the photoreceptor/bipolar cell synapse has also been observed in a model of photocoagulation of rabbit retina, where the laser ablation acutely removes a patch of photoreceptors while leaving the inner retina intact (Beier et al., 2017). After some days, nearby photoreceptors slowly migrate toward and fill in the lesioned area (Sher et al., 2013). As they do so, they form functional contacts with the deafferented bipolar cells (Sher et al., 2013; Beier et al., 2017). Another example of plasticity at the photoreceptor/bipolar cell synapse is the AAV-mediated gene therapy to replace retinoschisin (RS1) in adult mice (Ou et al., 2015). Retinal development of the RS1 knockout mice appears to proceed normally. However, the absence of RS1, a cell adhesion protein, eventually causes splitting of the retina and a failure of synaptic maintenance that manifests in reduction of the ERG b-wave amplitude (Sikkink et al., 2007). This defect was reversed upon RS1 gene replacement (Ou et al., 2015). Here we demonstrate for the first time that activation of rod input in young adults reversed synaptic changes that occurred during development and established functional contacts with their downstream neurons in the retinal circuitry of the adult retina. These results support the therapeutic potential of repairing or replacing defective rods in the degenerating retina. However, a critical time window for rescue likely exists: recent clinical trials for Leber Congenital Amaurosis to replace RPE65 in human patients for treating a type of Leber’s congenital amaurosis (LCA) caused by RPE65 mutations show limited success in visual improvement, and the retina continued to degenerate in some patients (Cideciyan et al., 2013). Future experimentation will address whether a critical time window of rescue exists for these approaches.

## Acknowledgements

This work was supported by National Institute of Health grants EY027193 (APS, GDF, and JC); EY12155 and EY027387 (JC); an unrestricted grant from Research to Prevent Blindness to the Department of Ophthalmology, UCLA; and Jules Stein Eye Institute Core Grant EY00331 (APS). We thank Dr. M. Scalabrino for comments on the manuscript, Dr. K.Martemyanov for providing the mGluR6 antibody, Dr. C. Craft for providing the cone arrestin (ARR3) antibody and Dr. S. Ruffins at the USC microscopy core for his help with confocal imaging.

